# Mutational analysis of a conserved positive charge in the *c*-ring of *E. coli* ATP synthase

**DOI:** 10.1101/2022.07.28.501891

**Authors:** Rashmi K. Shrestha, Michael W. Founds, Sam J. Shepard, Mallory M. Rothrock, Amy E. Defnet, P. Ryan Steed

## Abstract

F_1_F_o_ ATP synthase is a ubiquitous molecular motor that utilizes a rotary mechanism to synthesize adenosine triphosphate (ATP), the fundamental energy currency of life. The membrane-embedded F_o_ motor converts the electrochemical gradient of protons into rotation, which is then used to drive the conformational changes in the soluble F_1_ motor that catalyze ATP synthesis. In *E. coli*, the F_o_ motor is composed of a *c*_10_ ring (rotor) alongside subunit *a* (stator), which together provide two aqueous half channels that facilitate proton translocation. Previous work has suggested that Arg50 and Thr51 on the cytoplasmic side of each subunit *c* are involved in the proton translocation process, and positive charge is conserved in this region of subunit *c*. To investigate the role of these residues and the chemical requirements for activity at these positions, we generated eleven substitution mutants and assayed their *in vitro* ATP synthesis, H^+^ pumping, and passive H^+^ permeability activities, as well as the ability of mutants to carry out oxidative phosphorylation *in vivo*. While polar and hydrophobic mutations were generally tolerated in either position, introduction of negative charge caused a substantial defect. We discuss the possible effects of altered electrostatics on the interaction between the rotor and stator, water structure in the aqueous channel, and interaction of the rotor with phospholipids.

## 1. Introduction

F_1_F_o_ ATP synthase uses an electrochemical gradient of H^+^ or Na^+^ ions to complete the ultimate step in oxidative phosphorylation, making it the primary producer of ATP across all domains of life [1]. Due to its critical role in cellular bioenergetics and potential role in mitochondrial dysfunction and cell death [2,3], ATP synthase is an emerging drug target [4], with particular interest in novel antibiotics that inhibit ATP synthase of multidrug-resistant bacteria [5,6]. Structurally, ATP synthases are a complex of two rotary motors, F_o_ and F_1_. The ion-translocating F_o_ motor, composed of a rotor ring of 8-15 *c* subunits associated with a stator and peripheral stalk of varying composition, is embedded in the cell membrane of prokaryotes, the inner membrane of mitochondria, or the thylakoid membrane of chloroplasts. The F_1_ motor, which is extrinsic to the membrane, contains the active sites for ATP synthesis/hydrolysis and is composed of an α_3_β_3_ hexamer surrounding a γε central stalk and capped with a subunit that links F_1_ to the peripheral stalk. In mitochondria, F_1_F_o_ complexes dimerize via accessory proteins in F_o_ [7]. During ATP synthesis, ion translocation through F_o_ from the positive (P) side of the membrane to the negative (N) side generates rotation of the *c*-ring that is transmitted to the γ stalk of F_1_ to drive the conformational changes of the rotary binding change mechanism [8,9]. In some cases, ATP synthase can operate in reverse, hydrolyzing ATP to pump H^+^ against the electrochemical gradient. Bacterial ATP synthases are monomeric and have the simplest subunit composition.

The structure of *E. coli* ATP synthase has been captured in several rotational states by cryo-electron microscopy (cryo-EM) at sufficient resolution to identify key residues in the F_o_ motor (Fig. 1) [10,11]. Stator subunit *a* is composed of five transmembrane (TM) helices and sits between the subunit *b* dimer, which forms the peripheral stalk, and the decameric *c*-ring, where each *c* subunit is a hairpin of two TM helices. TM helices 2-5 of subunit *a* are nearly horizontal relative to the membrane plane and abut the surface of the *c*-ring to form an extensive interacting surface centered around the essential Arg210 residue in TM helix 4 of subunit *a* and the H^+^-binding Asp61 residue in TM helix 2 of each *c* subunit. Two offset aqueous half channels that make up the H^+^ translocation pathway through subunits *a* and *c* have been mapped by probing reactivity of substituted cysteine residues [12] and are confirmed by the cryo-EM structure [11]. The periplasmic (P-side) half channel extends through the middle of the TM 2-5 four-helix bundle from the periplasmic surface up to Arg210, and the cytoplasmic (N-side) half channel lies at the interface between subunit *a* and the *c*-ring extending from Asp61 to the cytoplasmic surface. During gradient-driven ATP synthesis, these half channels facilitate the protonation and deprotonation of Asp61 residues on the *c*-ring resulting in rotation.

**Figure 1:**
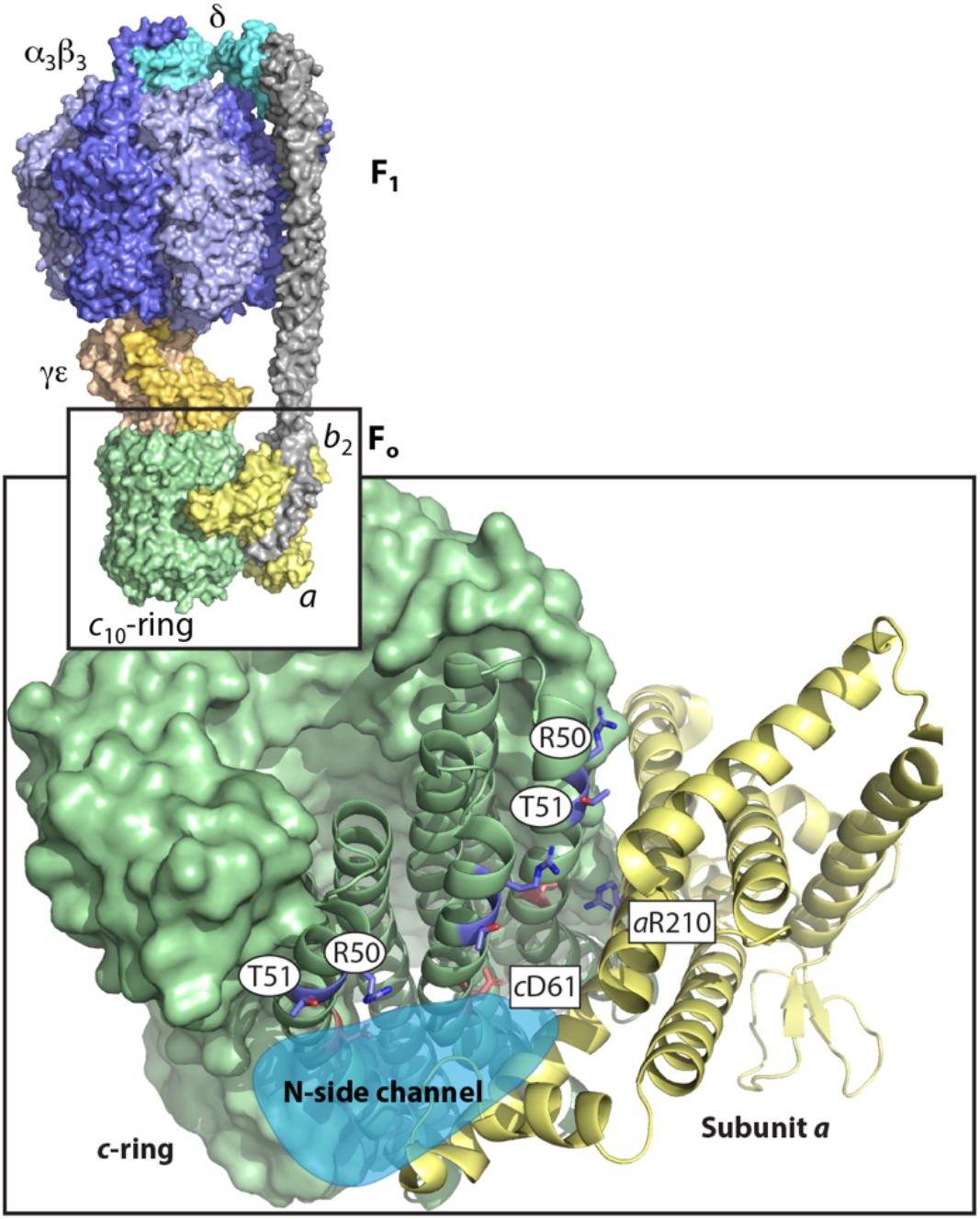
Structural context of Arg50 and Thr51. Structure of *E. coli* ATP synthase (PDB 6OQW) with detail showing the *a-c* interface of F° (PDB 6VWK) as viewed from the cytoplasmic (N) side. Essential *a*Arg210 and *c*Asp61 residues are marked for context, and the approximate extent of the N-side proton channel is shown in light blue.

While the functions of some critical residues along the H^+^ translocation pathway have been explored [13–17], questions remain about several polar or charged residues in the N-side half channel, including Arg50 and Thr51 on the cytoplasmic end of TM helix 2 of subunit *c*. Previous studies have noted that positive charge is conserved in this location across bacterial, mitochondrial, and chloroplastic ATP synthases, though the exact position varies by a helical turn [18,19]. Additionally, the previous Cys reactivity scan indicated that Cys substitution at Arg50 and the neighboring Thr51 disrupted ATP-driven H^+^ pumping activity [20]. To examine the roles of these residues in the proton translocation mechanism, we characterized six substitutions of Arg50 and five substitutions of Thr51, including non-polar, polar, and charged residues to examine the impact of these chemical properties on the functions of ATP synthase.

## 2. Results

### 2.1. Substitutions of Arg50

We constructed six substitution mutations to probe the function of Arg50, which lies on the surface of the *c*-ring near the cytoplasmic side of the membrane and forms part of the N-side H^+^ channel when interacting with subunit *a* (Fig. 1). Given the conservation of positive charge in this region [18], we tested the hypothesis that the positive charge on Arg is important for function. Mutations to Lys and His tested whether other positively charged residues could fulfill the role of Arg. Mutation to Ala or Met neutralized the charge and either removed or retained the steric bulk of the side chain. Finally, mutation to Glu or Asp reversed the charge at this position. These mutations were made by replacing the *uncE* gene in the whole-operon plasmid pFV2 [21] with a mutant gene fragment produced by PCR [22] or synthesized *de novo* (Twist Bioscience). Mutations were verified by Sanger sequencing prior to transformation of the *unc* operon deletion strain DK8 [23] for biochemical testing.

### 2.2. Negative charge in place of Arg50 inhibited H^+^ pumping and ATP synthesis

Inverted membrane vesicles prepared from mutant *E. coli* strains were assayed for ATP-driven H^+^ pumping and H^+^-driven ATP synthesis activities as described in Section 4, and results are summarized in Figure 2. His and Lys substitutions retained function in ATP synthesis and H^+^ pumping. Surprisingly, non-polar substitutions were generally tolerated at position 50 with only mild inhibition of H^+^ pumping activity. However, introduction of negative charge was detrimental to both ATP synthesis and H^+^ pumping activity, where Arg50Asp caused a much more significant defect than Arg50Glu.

**Figure 2:**
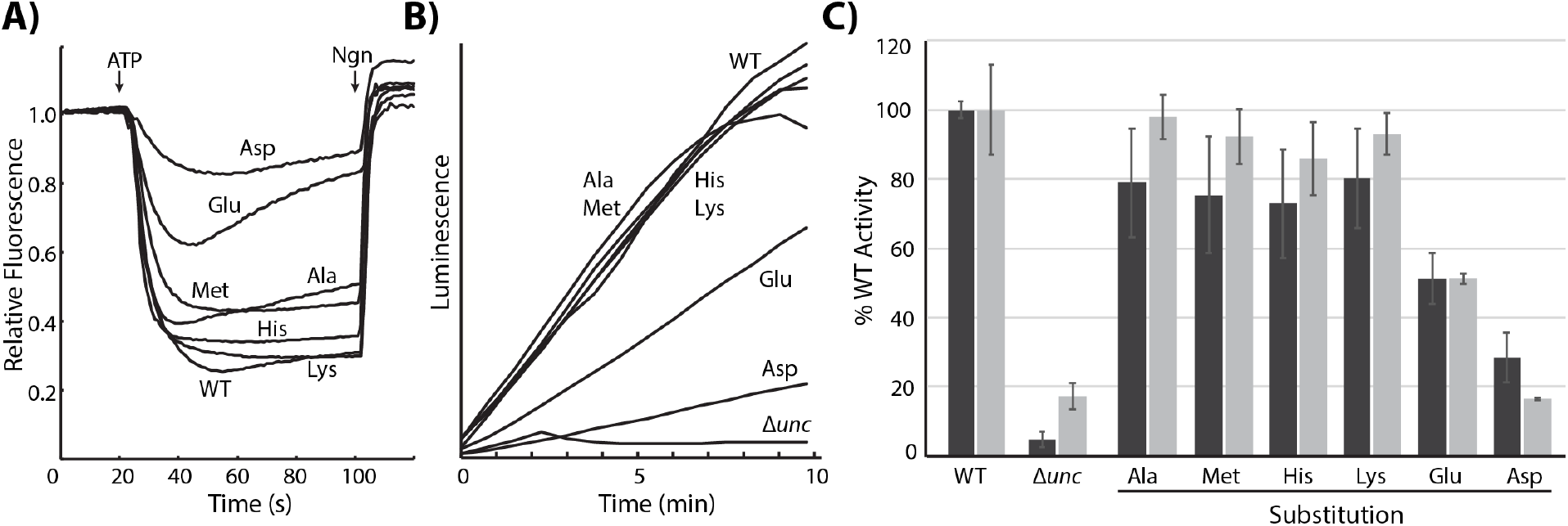
H^+^ pumping and ATP synthesis activities of Arg50 mutants. **A)** Representative ACMA fluorescence time traces show ATP-driven H^+^ pumping activity. Markers indicate the addition of ATP at 20s and the addition of nigericin (Ngn) at 100s. Fluorescence values for each trace were normalized to the initial value. **B)** Representative luminescence time traces are shown for H^+^ gradient-driven ATP synthesis activity. Inverted membrane vesicles were energized with NADH at *t*=0, and luminescence (arbitrary units) was monitored for 10 min. **C)** Activities of Arg50 mutants are plotted relative to WT. ATP synthesis activity (light bars) is the maximum slope (luminescence/s) defined by five consecutive points. Pumping activity (dark bars) is the percent fluorescence quenching (see Section 4.3). Activities are plotted as mean ± standard error (n≥3).

### 2.3. Substitutions of Thr51

We constructed five substitution mutations to probe the function of neighboring Thr51 (Fig. 1). Given that mutation to Cys, previously reported as non-functional [20], would remove polarity and hydrogen bonding potential, we prepared mutations to test the hypothesis that these properties are important for the function of Thr51. Mutation to Ser retained the hydroxyl while increasing polarity and reducing steric bulk. Mutation to Ala, Val, and Leu eliminated the hydroxyl and tested side chains smaller than, isosteric with, and larger than Thr, respectively. Finally, mutation to Asp tested a further increase in polarity and introduced negative charge density to the surface of the *c*-ring.

### 2.4. Negative charge in place of Thr51 inhibited H^+^ pumping and ATP synthesis

Inverted membrane vesicles prepared from mutant *E. coli* strains were assayed for ATP-driven H^+^ pumping and H^+^-driven ATP synthesis activities as above, and the results are summarized in Figure 3. Most mutations caused mild inhibition of H^+^ pumping and ATP synthesis, and only Thr51Asp significantly disrupted both activities.

**Figure 3:**
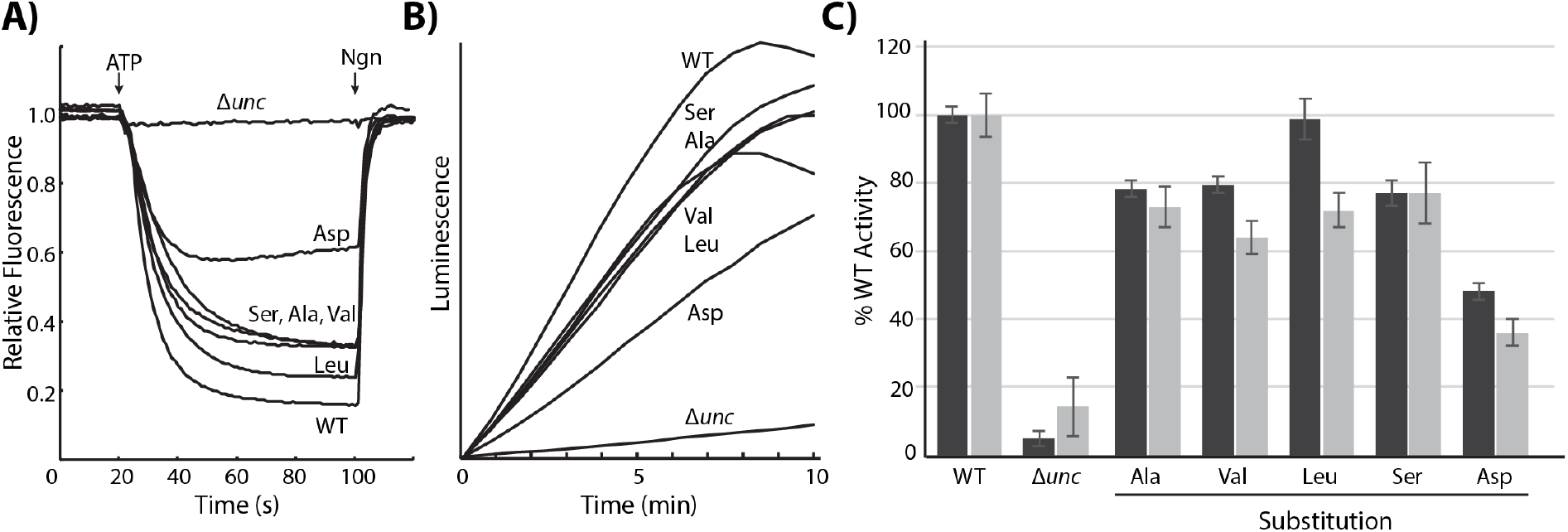
H^+^ pumping and ATP synthesis activities of Thr51 mutants. **A)** Representative ACMA fluorescence time traces. Markers indicate the addition of ATP at 20s and the addition of nigericin (Ngn) at 100s. Fluorescence values for each trace were normalized to the initial value. **B)** Representative luminescence time traces are shown for H^+^ gradient-driven ATP synthesis. Inverted membrane vesicles were energized with NADH at *t*=0, and luminescence (arbitrary units) was monitored in real time for 10 min. **C)** Activities of Thr51 mutants are plotted relative to WT. Pumping activity (dark bars) is the percent fluorescence quenching (see Section 4.3). ATP synthesis activity (light bars) is the maximum slope (luminescence/s) defined by five consecutive points. Activities are plotted as mean ± standard error (n≥3).

### 2.5. Arg50Asp blocked passive H^+^ transport

To further characterize the effects of the above substitutions at Arg50 or Thr51 on the movement of H^+^ through F_o_, we measured the passive translocation of H^+^ through F_o_ after the F_1_ sector had been stripped from the membrane. Stripped membrane vesicles were energized with NADH, and gradient formation was monitored using ACMA fluorescence quenching. In contrast to the ATP-driven H^+^ pumping experiments above, fluorescence quenching would indicate a blockage of H^+^ movement through F_o_ and therefore a disruption of normal function. In general, the effects of substitutions on passive permeability reflected those on H^+^ pumping and ATP synthesis activities. Arg50 substitutions fell into three categories, where Arg50Asp caused a complete blockage in H^+^ translocation, Arg50Ala and Arg50Glu were moderately blocked, and the other substitutions only slightly reduced H^+^ translocation relative to WT (Fig. 4, A). All of the Thr51 substitutions caused only minor disruption, if any, relative to WT (Fig. 4, B).

**Figure 4:**
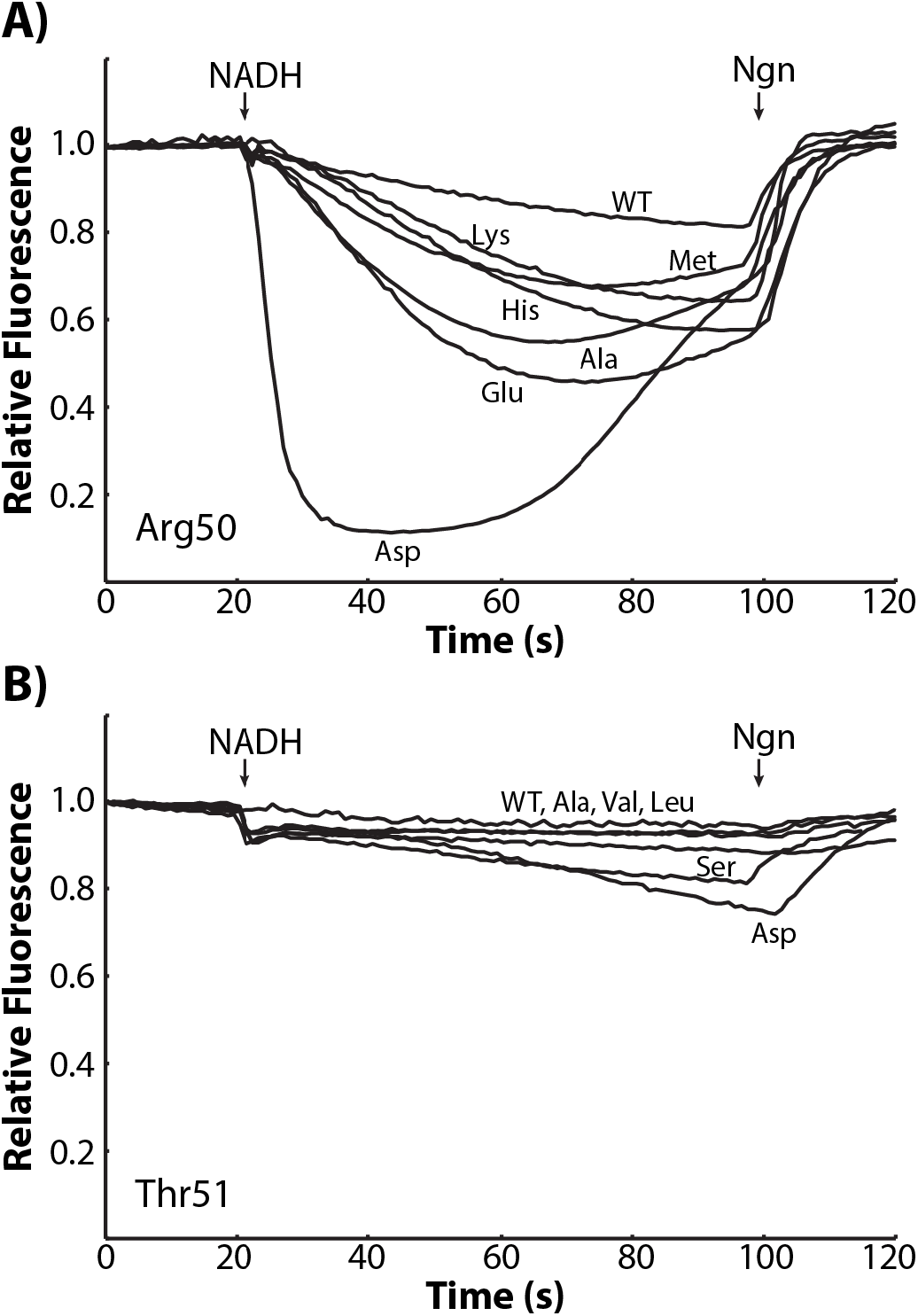
Effects of substitutions on passive H^+^ translocation. Representative ACMA fluorescence time traces are shown for stripped membrane vesicles of Arg50 substitutions **(A)** and Thr51 substitutions **(B)**. Markers indicate the addition of NADH at 20 s and the addition of nigericin (Ngn) at 100 s. Fluorescence values for each trace were normalized to the initial value.

### 2.6. Only Arg50Asp prevented growth of E. coli on succinate

To determine whether the above mutations affected the activity of ATP synthase *in vivo*, we measured the growth of *E. coli* mutants on succinate minimal medium, where succinate is the sole carbon source and growth requires a functional oxidative phosphorylation system. Consistent with other characterizations, most substitutions at Arg50 and Thr51 caused only a mild growth defect (Fig. 5). However, Arg50Ala caused a moderate growth defect, and growth of the Arg50Asp mutant was indistinguishable from the DK8 (Δ*unc*) negative control.

**Figure 5:**
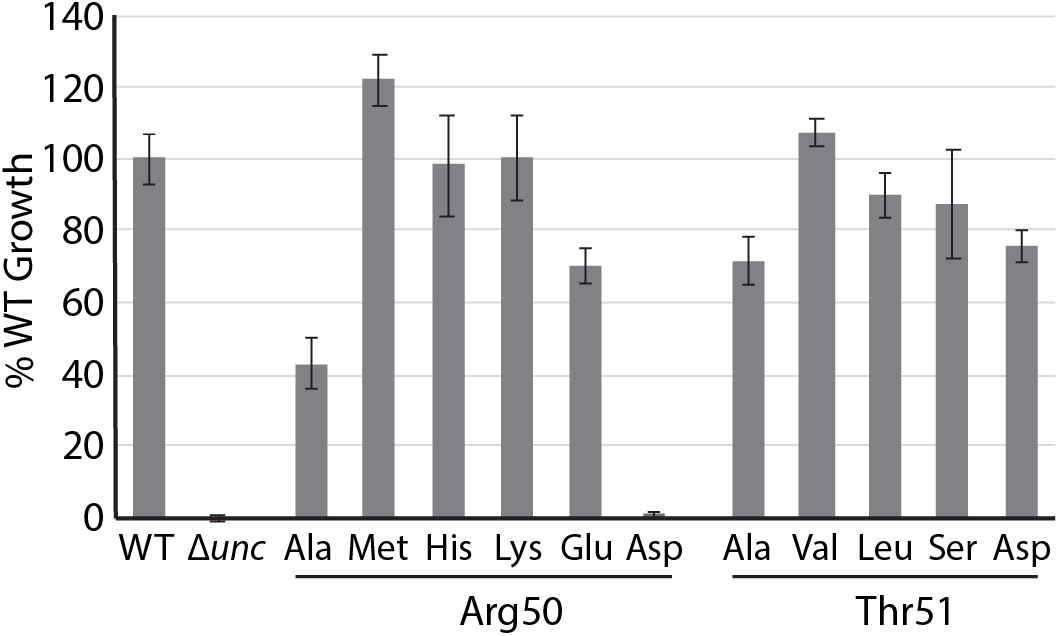
Effects of mutations on succinate growth. *E. coli* mutants were grown on M63-TIV minimal medium supplemented with glucose or succinate (see Section 4.6) and cell density was monitored by OD_550_ over 8 hours. Maximum cell density during the growth period was normalized to WT. Bars represent mean ± standard error (n≥3) relative to WT.

### 2.7. Substitutions did not prevent membrane insertion and assembly

We confirmed that none of the mutations prevented the membrane insertion and assembly of subunit *c* by detecting the presence of subunit *a* in membrane vesicles by Western blotting (Fig. 6). Since the insertion of subunit *a* requires the presence of subunits *b* and *c* in the membrane [24] and free subunit *a* is degraded unless assembled into F_o_ [25], detection of subunit *a* in the membrane reports proper assembly of the *c*-ring and the F_o_ complex. Therefore, detection of subunit *a* in inverted membrane vesicles from mutants, including Arg50Asp, indicated that observed functional defects arise from the alteration of the side chain rather than a failure of the F_o_ complex to assemble.

**Figure 6:**
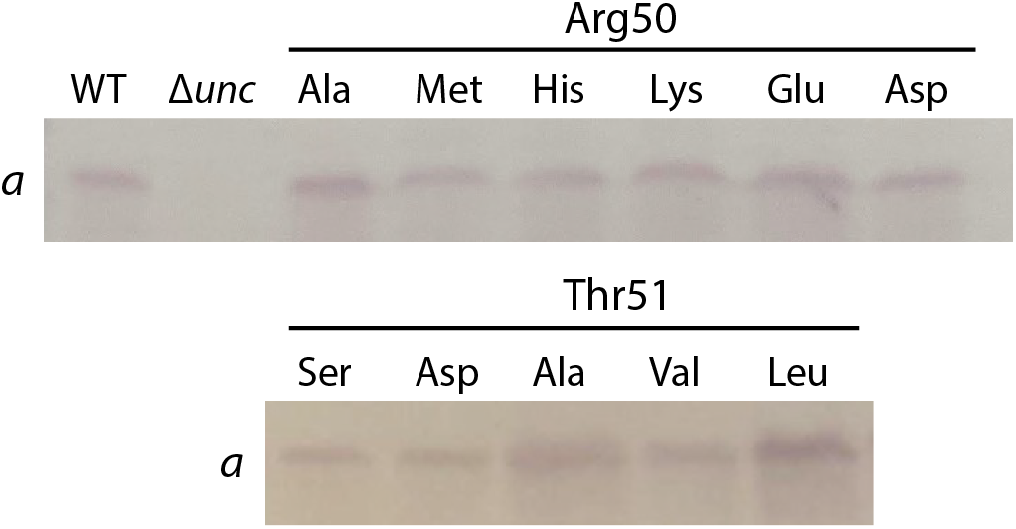
No effect of mutations on insertion of F_o_ subunits. Western blots of WT and mutant inverted membrane vesicles using antisera against subunit *a*. DK8 (Δ*unc*) was included as a negative control.

## 3. Discussion

We have demonstrated that a positive charge on the cytoplasmic side of TM helix 2 of subunit *c*, though conserved in this region, is not strictly necessary for H^+^ pumping or ATP synthesis activities. Indeed, substitutions of Arg50 and the neighboring Thr51 that neutralize the charge or polarity and change the size of the side chains are generally well tolerated. However, introducing negative charge density into either of these positions reduces both ATP synthesis and H^+^ pumping activities and blocks passive H^+^ translocation. The results show that changes in the electrostatics at these positions in the *a-c* interface can have significant effects on H^+^ translocation.

Positive charge is conserved on the N-side of TM helix 2 of subunit *c* [18] as are small polar side chains like Thr, although their placement varies by one helical turn (Fig.7, A). In higher organisms, the conserved positive charge on the N-side of subunit *c* is a Lys, and there is evidence that the ε amino group is trimethylated [26]. The effects observed when these residues are mutated could arise from disruption of 1) a rotor-stator interaction between subunit *a* and subunit *c*, 2) water structure in the N-side aqueous half channel, or 3) a lipid interaction with the *c*-ring surface. Moreover, interpretation is complicated by the existence of *c*-ring surface residues at both the protein-lipid interface and the subunit *a-c* interface, and substitution mutations alter both interfaces.

**Figure 7:**
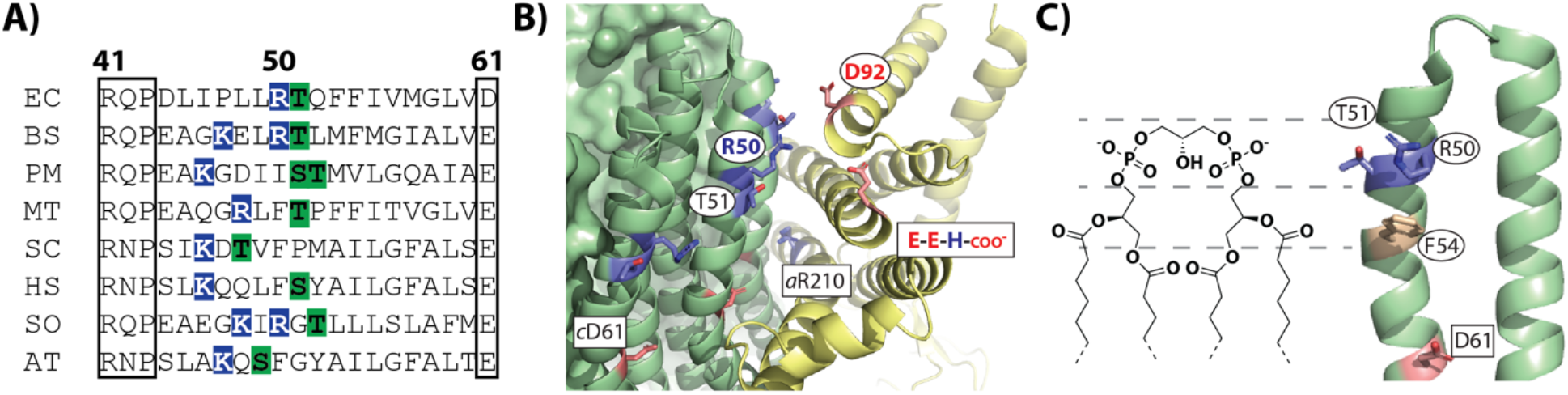
Conservation and possible interactions of Arg50 and Thr51. **A)** Amino acid sequence alignment of subunit *c* from several species in the region between the conserved R(Q/N)P loop motif and H^+^ binding acidic residue (open boxes). Arg and Lys residues in this region are highlighted in blue and nearby Ser and Thr residues are highlighted in green. EC, *Escherichia coli*; BS, *Bacillus subtilis*; PM, *Propionigenium modestum*; MT, *Mycobacterium tuberculosis*; SC, *Saccharomyces cerevisiae*; HS, *Homo sapiens*; SO, *Spinacia oleracea*; AT, *Arabidopsis thaliana*. **B)** Structural detail of the *a-c* interface (PDB 6VWK) showing proximity of Arg50 to negative charges in subunit *a*, including Asp92 and the acidic C-terminal residues (E-E-H). Note that the highlighted C-terminal residue (stick representation) is Glu269, as Glu270 and His271 are not included in the model. **C)** Approximate placement of a cardiolipin headgroup next to a single *c* subunit (PDB 6VWK) showing potential interactions consistent with the cardiolipin binding site proposed by Corey *et al*. [27].

### 3.1. Possible disruption of a rotor-stator interaction

Martin *et al*. proposed that Arg50 in *E. coli*, and equivalent Lys or Arg residues in other species, may form a transient electrostatic interaction with subunit *a* that enhances the H^+^-driven torque-generation mechanism during ATP synthesis [18]. Based on the most recent cryo-EM model [11], the nearest acidic residue to *c*Arg50 is *a*Asp92 in the cytoplasmic loop of subunit *a* between TM helices 1 and 2 (Fig. 7, B). In fact, Cys scanning indicated that *a*Asp92 contributes to the H^+^ translocation pathway [28] supporting a role for this residue in the H^+^ translocation mechanism. The C-terminus of subunit *a* is an additional source of negative charge density in nearly the same location (Fig. 7, B), including side chains of Glu269 and Glu270 and the terminal carboxylate of His271. Since removal of positive charge (Arg50Ala and Arg50Met) did not completely disrupt function, it seems that this putative electrostatic interaction is not strictly necessary for H^+^ pumping or ATP synthesis. On the other hand, introduction of like charges (Arg50Asp, Arg50Glu, or Thr51Asp) was disruptive, possibly by weakening the interaction between subunit *a* and the *c*-ring by electrostatic repulsion.

### 3.2. Possible disruption of hydrogen bonding

A second possibility is that alterations of polar and charged residues affect a hydrogen bonding network among *c* subunits and/or with water molecules in the N-side channel. In the cryo-EM model [11], the side chain guanidinium of Arg50 is close enough to form hydrogen bonds with Thr51 of the same *c* subunit or Gln52 of an adjacent *c* subunit. These inter-subunit interactions may contribute to maintaining the structural integrity and stability of the *c*-ring in the membrane. Since a substitution in the *uncE* gene will result in a substitution in all 10 copies of subunit *c* in the ring, one may expect even a small change in inter-subunit interactions to affect ring stability. However, perhaps surprisingly, we did not observe an assembly defect for any of the mutations tested.

For *c* subunits at the *a-c* interface, the side chains of both Arg50 and Thr51 extend into the cytoplasmic (N-side) aqueous half channel. Given the restricted space in this channel it is likely that water molecules do not behave as bulk water and are oriented by the polar (and non-polar) side chains lining the channel [29,30]. Others have proposed [31,32] that proton transport through the half channels occurs by a Grotthuss mechanism, which would require ordered water molecules in the aqueous half channels. Evidence for ordered water molecules is strongest for the P-side half channel [7,11], and while structured water molecules may exist deeper in the N-side channel, the tolerance of many substitutions at Arg50 and Thr51 argues against an exquisitely structured water wire at this point in the N-side channel where the pore would be widest. However, it is possible that introduction of negative charge along the surface of the *c*-ring would cause enough reorientation of water molecules to disrupt a Grotthuss mechanism in the N-side channel, thus leading to the observed functional defects.

### 3.3. Possible disruption of a rotor-lipid interaction

Finally, the results are consistent with the involvement of Arg50 and Thr51 in a binding site for phospholipids on the surface of the *c*-ring, particularly cardiolipin. Cardiolipin is found in bacterial cell membranes, as well as the inner membrane of mitochondria and the thylakoid membrane of chloroplasts, and is known to be important for oxidative phosphorylation, possibly by facilitating H^+^ binding and transport on the surface of the membrane [33]. It has been proposed that the trimethylated Lys present in subunit *c* of higher organisms is important for interactions between the *c*-ring and phospholipids, particularly cardiolipin, and that cardiolipin may enhance H^+^ release from the exit channel during synthesis [19]. Density consistent with cardiolipin has been discovered in proximity to this region in the cryo-EM structures of mitochondrial ATP synthases [7,34], as well as in *E. coli* ATP synthase [11], and a specific interaction between cardiolipin and *E. coli* subunit *c* has been detected by solid state NMR [35]. Corey *et al*. [27] posited a loosely defined cardiolipin binding motif, based on analysis of simulated cardiolipin binding to *E. coli* membrane proteins, in which binding is supported by Arg or Lys residues in the membrane plane, at least one polar residue (Ser or Thr), and at least one aromatic (or bulky hydrophobic) residue slightly deeper in the membrane. The Arg50 side chains from adjacent *c* subunits along with Thr51 and Phe54 residues resemble this cardiolipin binding motif (Fig. 7, C) in which the Arg positive charges would interact with the negatively charged phosphates, Thr51 would hydrogen bond with hydroxyls of the glycerol backbone, and Phe54 would interact with acylglycerols. Substitutions that neutralize the Arg charges may weaken but not abolish the selective association of lipids with the *c*-ring leading to the observed mild to moderate effects on activity that we observed. Likewise, substitutions with negative charge, especially in place of Arg50, would markedly increase the negative charge density in the region that likely interacts with negatively charged head groups and may lead to repulsion of particular lipids, including cardiolipin.

In summary, we have shown that Arg50 and Thr51 in subunit *c* tolerate most substitutions and are not strictly required for ATP synthesis or ATP-driven H^+^ pumping activity, despite the conservation of positive charge and polarity in this region. However, introduction of negative charge to either position does reduce both activities and blocks passive H^+^ transport. Whether these defects are caused by an alteration of a hydrogen bonding network or weakened interaction with either lipids or subunit *a* will be explored in future work.

## 4. Materials and Methods

### 4.1. Construction of substitution mutations

Mutations were introduced into the Cys-less plasmid construct pFV2 [21], which encodes all eight structural genes of ATP synthase and where all endogenous cysteine residues in F_1_ were substituted with alanine and the single cysteine residue in subunit *b* of F_o_ was substituted with serine. This construct is referred to as wild-type (WT). Either a two-step PCR method [22] or gene synthesis (Twist Bioscience) was used to generate mutant gene fragments that were then digested and ligated into cut pFV2 at the BsrGI and PpuMI restriction sites. Mutations were confirmed by DNA sequencing through the ligation sites, and the resulting mutant plasmids were transformed into the *unc* operon deletion strain DK8 [23], which does not produce functional ATP synthase.

### 4.2. Preparation of inverted membrane vesicles

DK8 transformant strains were grown in LB medium, and cells were harvested by centrifugation in the late exponential stage of growth. The pellets were resuspended in TMG buffer (50 mM Tris-acetate, 5 mM MgCl_2_, 10% (*v/v*) glycerol, pH 7.5) supplemented with 1 mM dithiothreitol and 1 mM phenylmethanesulfonyl fluoride. The suspension was disrupted by four passages through an Avestin EmulsiFlex-B15 homogenizer at ≥ 15000 psi. The lysate was centrifuged at 9000 × *g* for 10 min at 4 °C to remove intact cells and debris. Membrane vesicles were collected by centrifugation of the cleared lysate at 168,600 × *g* for one hour at 4 °C. Membrane pellets were washed once with TMG buffer and then resuspended in TMG buffer using a Dounce homogenizer and stored at -80 °C. Protein concentrations were determined using a modified Lowry assay [36].

### 4.3. Measurement of ATP-driven H^+^ pumping

Inverted membrane vesicles were diluted to 10 mg/mL using TMG buffer and 160 µl of the diluted vesicles were suspended in 3.2 mL HMK buffer (50 mm HEPES-KOH, 2 mM MgCl_2_, 300 mM KCl, pH 7.5) at room temperature, and 9-amino-6-chloro-2-methoxyacridine (ACMA) was added to 0.3 μg/mL. Fluorescence data was collected on a Shimadzu RF-6000 spectrofluorometer. Baseline fluorescence was recorded at an excitation wavelength of 415 nm and an emission wavelength of 485 nm. The fluorescence quenching was initiated by adding 30 μL of 25 mM ATP and terminated by adding nigericin to 0.5 μg/mL. The relative quenching of the membrane vesicles was calculated using the return fluorescence after the addition of nigericin.

### 4.4. Measurement of ATP synthesis

ATP synthesis was measured using a luciferin/luciferase assay adapted from Preiss *et al*. [37]. In a 96-well white microplate, wells contained 180 µL reaction mixture (5 mM Tricine, pH 8.0, 50 mM KCl, 2.5 mM MgCl_2_, 3.75 mM potassium phosphate, 0.1 mM adenosine diphosphate, 2.5 mM reduced β-nicotinamide adenine dinucleotide (NADH), 125 µM luciferin, and 100 ng luciferase). The synthesis reaction was initiated by mixing in 20 µL 0.5 mg/mL inverted membrane vesicles. Luminescence was measured every 20-30 s for 10 min at 30 °C using a BioTek Synergy H1 microplate reader. ATP synthesis activity (luminescence/min/mg) was defined as the maximal slope of five consecutive points during the assay.

### 4.5. Measurement of passive proton permeability

F_1_ was chemically stripped from the inverted membrane vesicles using a low salt, high pH buffer as previously described [20], and the protein concentration of stripped vesicle preparations was determined by a modified Lowry assay as above. Proton permeability was assayed using ACMA fluorescence quenching as described in Section 4.3. However, the amount of stripped inverted membrane vesicles used was 480 µg and fluorescence quenching was initiated by addition of 16 μL of 10 mM NADH.

### 4.6. Mutant growth on succinate minimal medium

*E. coli* DK8 transformant strains were streaked on an LB agar plate with 100 µg/mL ampicillin and single colonies were used to inoculate M63-TIV minimal medium (61.8 mM KH_2_PO_4_, 38.2 mM K_2_HPO_4_, 15 mM (NH_4_)_2_SO_4_, 1 mM MgSO_4_, 1 µg/mL thiamine, 0.2 mM isoleucine, 0.2 mM valine) containing 0.1% (*w/v*) glucose and 5% (*v/v*) LB. After overnight growth at 37 °C, 5 µL dense culture was used to inoculate wells in a clear 96-well microplate, each containing 150 µL M63-TIV minimal medium with 0.04% (*w/v*) glucose or 0.6% (*w/v*) succinate as a carbon source. All growth media contained 100 µg/mL ampicillin except when culturing untransformed DK8. Growth at 37 °C with shaking was monitored by measuring OD_550_ every 30 min for 12 hours.

### 4.7. Detection of subunit a by western blotting

Inverted membrane vesicles (50 µg protein) were run on a Bio-Rad AnyKD SDS-PAGE gel, and protein was subsequently transferred to a polyvinylidene fluoride membrane in Towbin buffer (25 mM Tris-HCl, 192 mM glycine, 20% methanol) at 70 V for 90 min. The membrane surface was blocked with 5% (*w/v)* dry milk in TBST buffer (20 mM Tris-HCl, pH 7.6, 150 mM sodium chloride, 0.1% (*v/v*) Tween-20). After rinsing in TBST, the membrane was immunostained with rabbit anti-subunit *a* antisera [15] (1:1000 in TBST with 2% (*w/v*) bovine serum albumin), rinsed with TBST, and then treated with alkaline phosphatase (AP)-conjugated goat anti-rabbit-IgG (Southern Biotech; 1:8000 in TBST with 2% (*w/v*) bovine serum albumin). After rinsing with TBST, labeled protein was visualized using the AP substrate 5-bromo-4-chloro-indolyl-phosphate/nitro blue tetrazolium (BCIP/NBT; Southern Biotech).

## Abbreviations used

ACMA: 9-amino-6-chloro-2-methoxyacridine
AP: alkaline phosphatase
ATP: adenosine-5’-triphosphate
EM: electron microscopy
HEPES: 4-(2-hydroxyethyl)-1-piperazineethanesulfonic acid
NADH: reduced β-nicotinamide adenine dinucleotide
PAGE: polyacrylamide gel electrophoresis
PDB: protein data bank
SDS: sodium dodecyl sulfate
TM: transmembrane
WT: wild-type

## Acknowledgements

The authors acknowledge preliminary work by Clara Ledsky, Rahul Uppal, and students in the Spring 2016 CHE321 lab at Carleton College, Northfield, MN. Research was supported by National Institutes of Health grant R15 GM134453 to PRS and by the North Carolina GlaxoSmithKline Foundation.

## Author Contributions

PRS, MWF, and RKS designed research; RKS, MWF, and SJS constructed mutants; RKS, MMR, SJS, AED, and PRS performed biochemical experiments; RKS and PRS wrote the manuscript; all authors edited the manuscript.

